# Systematic Design and Comparison of Expanded Carrier Screening Panels

**DOI:** 10.1101/080713

**Authors:** Kyle A. Beauchamp, Dale Muzzey, Kenny K. Wong, Gregory J. Hogan, Kambiz Karimi, Sophie I. Candille, Nikita Mehta, Rebecca Mar-Heyming, K. Eerik Kaseniit, H. Peter Kang, Eric A. Evans, James D. Goldberg, Gabriel A. Lazarin, Imran S. Haque

**Affiliations:** Counsyl. 180 Kimball Way, South San Francisco, CA 94080.

**Keywords:** Expanded Carrier Screening, Next Generation Sequencing

## Abstract

**Purpose:** The recent growth in pan-ethnic expanded carrier screening (ECS) has raised questions about how such panels might be designed and evaluated systematically. Design principles for ECS panels might improve clinical detection of at-risk couples and facilitate objective discussions of panel choice.

**Methods:** Guided by medical-society statements, we propose a method for the design of ECS panels that aims to maximize the aggregate and per-disease sensitivity and specificity across a range of Mendelian disorders considered serious by a systematic classification scheme. We evaluated this method retrospectively using results from 474,644 de-identified carrier screens. We then constructed several idealized panels to highlight strengths and limitations of different ECS methodologies.

**Results:** Based on modeled fetal risks for “severe” and “profound” diseases, a commercially available ECS panel (Counsyl) is expected to detect 183 affected conceptuses per 100,000 US births. A screen’s sensitivity is greatly impacted by two factors: (1) the methodology used (e.g., full-exon sequencing finds more affected conceptuses than targeted genotyping), and (2) the detection rate of the screen for diseases with high prevalence and complex molecular genetics (e.g., fragile X syndrome).

**Conclusion:** The described approaches allow principled, quantitative evaluation of which diseases and methodologies are appropriate for pan-ethnic expanded carrier screening.

## INTRODUCTION

Carrier screening attempts to identify couples at elevated risk of conceiving a pregnancy affected with a Mendelian condition ^1^, thereby enabling consideration of alternative reproductive options and early intervention strategies ^2,3,4^ There are thousands of Mendelian conditions ^1^ that differ in both incidence and severity, and for myriad reasons, carrier screening only interrogates a subset of these conditions. Indeed, current guidelines issued by the American Congress of Obstetricians and Gynecologists (ACOG) and American College of Medical Genetics and Genomics (ACMG) ^5^ suggest pan-ethnic screening only for cystic fibrosis (ACOG, ACMG) ^6,7^ and spinal muscular atrophy (ACMG). ^8^

Introduced in 2009, “expanded” carrier screening (ECS), which identifies reproductive risks for dozens to hundreds of diseases, has gained acceptance as a reasonable screening approach ^5 9^. ECS offerings are diverse, spanning a range of panel sizes and assay technologies. Two ostensibly identical ECS panels with the same number of genes and assay technology may nevertheless differ in their sensitivity due to differences in the number of interrogated positions in each gene and in the interpretation of detected variants. In principle, maximal sensitivity is achieved by determining the sequence at every base in an entire gene and curating all variants to assess pathogenicity. However, due to assay cost and throughput limitations in variant interpretation, the set of interrogated bases is typically limited. In the ECS strategy termed “full-exon sequencing”, most intronic bases are not included in the panel, and next-generation sequencing (NGS) is used to identify all remaining bases across the protein-coding exons and in noncoding regions with known contributions to pathogenesis (e.g., known splice-modifying sites). Full-exon sequencing typically probes thousands of bases per gene and can identify all common variants, plus rare novel variants such as clearly damaging protein-truncating mutations. Because of its potential to discover novel variants, achieving high sensitivity and specificity with full-exon sequencing requires processes to interpret the clinical impact of all observed variants. An alternative ECS strategy that sidesteps the need for novel-variant curation is “targeted genotyping” (TG), which restricts its focus to a set of predefined pathogenic variants, often between 1 and 50 per gene ^9^ (but in some cases, such as *CFTR* mutation panels, ranging into hundreds per gene). Though NGS can be used to perform TG, other technologies, such as allele-specific PCR or microarray, can also be used. Interestingly, even though TG-based tests may be implemented inexpensively (because of a smaller assayed region and no need for novel variant interpretation), their lower detection rate relative to full-exon sequencing may increase overall health-care spending due to the increased cost of care for undetected affected pregnancies. ^10^

Recently, several professional societies, including ACOG and ACMG, have provided recommendations regarding the broader implementation of ECS. Their suggestions include careful vetting of the clinical and population characteristics of each disease and selection of panel content using ^11^ “clear criteria, rather than simply including as many diseases as possible.” Similarly, the European Society for Human Genetics recommended that ^12^ “an important screening criterion is that the natural course of the disease screened for should be adequately understood, and that an acceptable and reliable test should be available with known sensitivity, specificity and predictive values.”

To address these points and increase the transparency of expanded carrier screening panel design, this study proposes a method for the systematic design of ECS tests. This approach builds upon previous work in three main ways: by focusing on diseases with high clinical significance ^2,13^, by using well-calibrated curations ^14^ to characterize variants, and by extending a framework suggested previously ^15^ for systematically characterizing the performance of expanded carrier screening panels. We evaluate and discuss this method using ECS data from more than 400,000 patients.

## MATERIALS AND METHODS

### Systematic Design of Expanded Carrier Screening

In this work, we propose the following approach to designing expanded carrier screening panels:

1. Systematically enumerate all candidate diseases for which universal screening is clinically desirable (typically “severe” or “profound” diseases, as defined below).
2. Maximize the aggregate panel sensitivity subject to limitations on assay size by selecting candidates that capture a high amount of disease risk (i.e., high-incidence diseases).
3. Maximize per-disease sensitivity and negative predictive value (NPV) to yield high confidence in non-carrier status for individual conditions.
4. Ensure near-100% specificity using carefully designed assay and curation protocols.

### Dataset Summary

Retrospective analysis of anonymized ECS data is exempt from Institutional Review Board oversight (as granted by Western IRB on 10/05/2016). All patients provided informed consent for testing and anonymized research. Data are based on de-identified, aggregated ECS results of 474,644 patients who were screened using the Family Prep Screen (Counsyl, South San Francisco, CA) for up to 94 “severe” or “profound” conditions (as described below). For congenital adrenal hyperplasia, we analyze results only for the classical form and not the nonclassical form, as the nonclassical form is classified as “moderate” severity (as described below). To minimize bias in disease frequencies, patient data were only used if the patient reported no remarkable personal or family history (e.g., infertility, or known history of carrier status or genetic disease). Using a method previously described, data from patients tested under two methodologies (344,743 targeted genotyping-based tests and 129,901 NGS-based tests) were aggregated to reduce statistical uncertainty ^15^. Statistical calculations were performed using Python 2.7.12, Numpy 1.11.1, and Pandas 0.18.1, as well as additional tools described previously ^15^. To assess the importance of panel-wide copy number variant (CNV) deletion calling, data from 56,267 NGS-based tests (primarily a subset of the 129,901 total NGS cohort) was analyzed to estimate the frequency of exon-or-larger deletions and duplications in 82 genes on the panel (method described in the Supplement). Deletions were assumed pathogenic, while duplications were excluded from further analysis. Unless otherwise noted, reported disease risks are weighted by the ethnic distribution of the United States ^16^, as described in the Supplementary Methods and Supplementary Table S1. Variants of uncertain significance (VUS) are not reported during routine carrier screening and are excluded from the present retrospective analysis (but are included in our assessment of curation in Table 3). Furthermore, variants with known low penetrance or mild phenotype have been excluded from analysis, as described previously (Table S6 in reference 15). After exclusion of low-penetrance and mild variants, all variants are treated as having equal phenotypic impact (excluding Fragile X syndrome, described below). As a case study, we dissect the most prevalent pathogenic variants in GBA in Table S5.

### Disease Selection by Clinical Severity

An ECS best serves patients by screening for serious diseases.^12^ Because the classification of large disease lists requires substantial effort on the part of medical professionals, we previously developed ^2^ a rule-based scheme that classifies diseases into increasingly serious categories (“mild”, “moderate”, “severe” or “profound”), using disease characteristics (e.g., intellectual disability or shortened lifespan) as inputs. Under this scheme, disease characteristics are grouped into four tiers, with Tier 1 being the most serious. This phenotype-based scheme was shown to agree with severity classifications by health-care professionals ^2^, suggesting its feasibility for efficient classification and comparison of hundreds of diseases. Underscoring the usefulness of such severity classifications, a separate survey of at-risk couples with “severe” or “profound” diseases found that among at-risk couples (those found to both be carriers for the same autosomal recessive condition), those at-risk for severe or profound conditions altered reproductive decisions at a significantly higher rate than those carrying moderate conditions ^13^.

In the present work, we propose that the first step in ECS disease selection is to enumerate and prioritize diseases considered “severe” or “profound” under this scheme. Although they may not affect reproductive choices, “moderate” conditions (such as *GJB2*-related hearing loss) may also be candidates for enumeration due to strong clinical desire for testing by patients and providers to enable early preparation. “Mild” conditions are typically screened in an as-requested (rather than routine) fashion. Note that disease characterization is performed at the level of diseases, rather than variants.

### Quantifying ECS Sensitivity

In our proposed ECS design methodology, diseases are selected in order to maximize the aggregate sensitivity among “severe” and “profound” diseases. Intuitively, this is achieved by selecting diseases with high prevalence; however, the relationship between prevalence and sensitivity must be made precise.

Historically, in the context of ECS panels, disease frequency has often been discussed in terms of carrier frequency and at-risk couple frequency ^17 18^. However, these metrics have limitations that prevent their use in assessing sensitivity. Carrier frequency is a suboptimal proxy because a single carrier result alone is not clinically actionable: the reproductive risk is a function of both parents for autosomal recessive (AR) diseases. The at-risk couple frequency—frequently the square of carrier frequency for autosomal conditions—is problematic because certain single-gene diseases have complex inheritance patterns that may modulate the risk of transmission to offspring. In fragile X syndrome, for example, fetal risk is not easily derived from the carrier frequency and instead requires a risk model that considers the probability of repeat expansion as a function of maternal CGG repeat number ^19^. For these reasons, the “modeled fetal disease risk”, defined as the disease probability in a conceptus of randomly selected male and female parents, was recently introduced by Haque and coauthors to quantify the relative yields of different screening panels ^15^. A complete mathematical and computational definition of disease risk calculation is given in the supplementary methods, with disease-specific details described previously (Supporting Information in reference 15). Modeled fetal disease risk statistically quantifies the rate of affected conceptuses using simulated parental populations and accounts for the various transmission rates of autosomal-recessive, X-linked, and complex (e.g., fragile X syndrome) diseases.

Here we define sensitivity as the fraction of disease risk, summed over all diseases on the panel, that an ECS panel is able to detect:

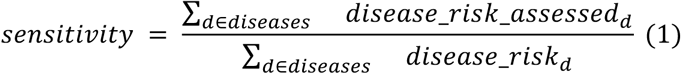

Here “disease_risk_assessed” is the disease risk captured by a particular panel, while “disease_risk” refers to the “true” disease risk of the population. If a test for a simple autosomal recessive condition were to detect 50% of carriers, the sensitivity would be *50%* * *50% = 25%,* where the squaring occurs because affected fetuses inherit two damaged copies of a gene. In this case, the test only captures one quarter of the disease risk for that disorder and compromises aggregate sensitivity.

In the present work, the candidate panel disease list is fixed and so the denominator in equation 1 is constant, i.e., it is the population disease risk aggregated over 94 “severe” and “profound” diseases. Therefore, to compare sensitivities of different ECS strategies, we focus on the numerator, i.e., the assessed disease risk.

### Clinical Accessibility of Variants: Curation

Genomics assays, particularly those based on NGS, require careful processes for deciding which variants are clinically significant and must be reported to patients. This variant interpretation process directly impacts both sensitivity and specificity. While ACMG and the Association for Molecular Pathology (AMP) have established general recommendations for this process ^14^, the optimal interpretation process will depend on the particular disease and subsequent clinical actions being considered ^20^. In particular, there exists a tradeoff among labor, sensitivity, and specificity in interpretive processes. While there exist processes involving little labor (e.g., those relying simply on computational methods), it is difficult to achieve both high sensitivity and high specificity simultaneously in a low-labor process.

Figure S1 describes the interpretation process developed at Counsyl to serve the needs of a high-throughput population-screening laboratory. Interpretation in this pipeline occurs in “real time”, whereby any novel variant observed in a patient sample (i.e., one that has not otherwise been classified recently) undergoes curation using the above process prior to the release of the patient’s report. The protocol uses an automated pipeline to collect several lines of evidence (population frequency, *in silico* protein structure predictors, splicing predictors, and conservation) that are then manually reviewed in combination with published case reports and functional studies to determine the variant classification. Prior to manual review, a rule-based system is used to automatically classify variants with high frequency in asymptomatic populations and variants with no literature reports.

To evaluate the variant classification performance of this pipeline, we compared Counsyl Family Prep Screen classifications to ClinVar ^21^ (April 2016 Release). As a reference standard, we selected variants that met the following requirements:

1. Each variant was classified by Counsyl and at least 2 of the following 9 external clinical labs: ARUP Laboratories, Clinical Biochemistry Laboratory University College London Hospitals, Emory Genetics Laboratory, GeneDx, Genetic Services Laboratory University of Chicago, Invitae, Juha Muilu Group Institute for Molecular Medicine Finland, LabCorp, and Laboratory for Molecular Medicine (Partners HealthCare). These laboratories were chosen due to having public classifications of genes and variants that sufficiently overlap with those on the Counsyl Family Prep Screen.
2. Each variant has complete concordance among classifications submitted by external laboratories.
3. Cancer susceptibility classifications for *ATM* and *NBN* were excluded due to lesser relevance for ECS applications.

The classifications were binned into two simplified categories according to medical management in carrier screening: (1) “Pathogenic”, which includes known pathogenic, likely pathogenic, and predicted pathogenic classifications, and (2) “VUS/benign”, which includes variant of uncertain significance (VUS), known benign, likely benign, and predicted benign. This process identified 505 variants located in 64 genes, of which 179 are pathogenic and 326 are VUS/benign. The molecular impact of the variants was obtained from the Ensembl variant effect predictor (VEP) web interface for human GRCh37 ^22^.

To demonstrate the tradeoff among labor, sensitivity, and specificity, we compare the performance of the Variant Effect Predictor ^22^ algorithm (VEP, a widely-used automated annotation method for sequence variants) under various sensitivity/specificity thresholds to the performance of a human-in-the-loop interpretive process. Specifically, we compare the Counsyl interpretive pipeline to “VEP Specific”, which prioritizes specificity, by flagging any variant having “High” VEP impact as pathogenic, and “VEP Sensitive”, which prioritizes sensitivity by flagging as pathogenic any variant having “High” or “Moderate” impact by VEP.

## RESULTS

### Compilation and classification of candidate genes

The first step in ECS panel design is the selection and severity-classification of a superset of candidate diseases. We performed a preliminary evaluation of 671 diseases and complete evaluation of 110 diseases—94 “Severe” and “Profound” conditions used in fetal disease risk calculations and 16 additional conditions ^23^. As described in Methods, diseases were classified by severity using objective criteria: as examples, cystic fibrosis and spinal muscular atrophy, recommended for pan-ethnic screening by ACMG ^24^, are classified as “severe”, whereas *GJB2*-related DFNB1 nonsyndromic hearing loss and deafness is classified as “moderate”. Severity classification of select diseases is shown in Table 1, with additional diseases given in Supplementary Table S2.

**Table 1.** The disease severity classification and phenotypic features are given for representative diseases. The severe and profound diseases were selected as commonly occurring on carrier screening panels. The fourth and least severe phenotypic feature group (Tier 4) contains only reduced fertility and is not shown.

| Disease Name | Gene | Severity | Phenotypic Features:<br>Tier 1 | Phenotypic Features:<br>Tier 2 | Phenotypic Features: Tier 3 | OMIM |
| --- | --- | --- | --- | --- | --- | --- |
| Smith-Lemli-Opitz syndrome | *DHCR7* | Profound | shortened life span:<br>infancy<br>intellectual disability | impaired mobility<br>internal physical<br>malformation | mental illness<br>dysmorphic features | #270400 |
| Carnitine palmitoyltransferase II deficiency | *CPT2* | Severe | shortened life span:<br>infancy<br>intellectual disability | impaired mobility<br>internal physical<br>malformation | sensory impairment<br>dysmorphic features | #608836 |
| Cystic fibrosis | *CFTR* | Severe | shortened life span:<br>childhood/adolescence | impaired mobility<br>internal physical<br>malformation | Immunodeficiency / cancer | #219700 |
| Fragile X syndrome | *FMR1* | Severe | intellectual disability | impaired mobility | sensory impairment: vision<br>immunodeficiency / cancer<br>mental illness<br>dysmorphic features | #300624 |
| Hb Beta Chain-Related Hemoglobinopathy | *HBB* | Severe | None | shortened life span:<br>premature adulthood<br>impaired mobility<br>internal physical<br>malformation | Immunodeficiency / cancer<br>sensory impairment: other | #603903 |
| Phenylalanine hydroxylase deficiency | *PAH* | Severe | intellectual disability | impaired mobility | None | #261600 |
| Spinal muscular atrophy | *SMA* | Severe | shortened life span:<br>infancy | impaired mobility | sensory impairment: touch,<br>other | #253300 |
| GJB2-related DFNB1 Nonsyndromic Hearing Loss and Deafness | *GJB2* | Moderate | None | None | sensory impairment: hearing | #220290 |
| Pseudocholinesterase deficiency | *BCHE* | Mild | None | None | None | *177400 |

### Disease Risk Comparison of Idealized ECS Panels

The next steps in our ECS panel-design framework involve maximizing aggregate and gene-level sensitivities. To identify the key factors that influence this maximization, we mined ECS data from 474,644 patients and reweighted the computed risks by the ethnic makeup of the United States ^16^. The per-gene contribution to the aggregate disease risk assessed by an ECS is shown in Figure 1. It is clear that a few well-known, high-prevalence diseases contribute substantially to the overall disease risk. Furthermore, several of these large contributors, such as *FMR1* (fragile X syndrome, 15.8% of total disease risk) and *CYP21A2* (CAH; 21-hydroxylase deficient congenital adrenal hyperplasia, 5.2% of total disease risk), arise from genes requiring special care in analysis (Table 2; see Discussion). Over one-fourth (28.9%) of total disease risk is accountable to these four technically challenging conditions, suggesting that in order for an ECS to maximize assessment of disease risk, sensitivity for technically challenging yet highly prevalent diseases is critical.

**Figure 1.**
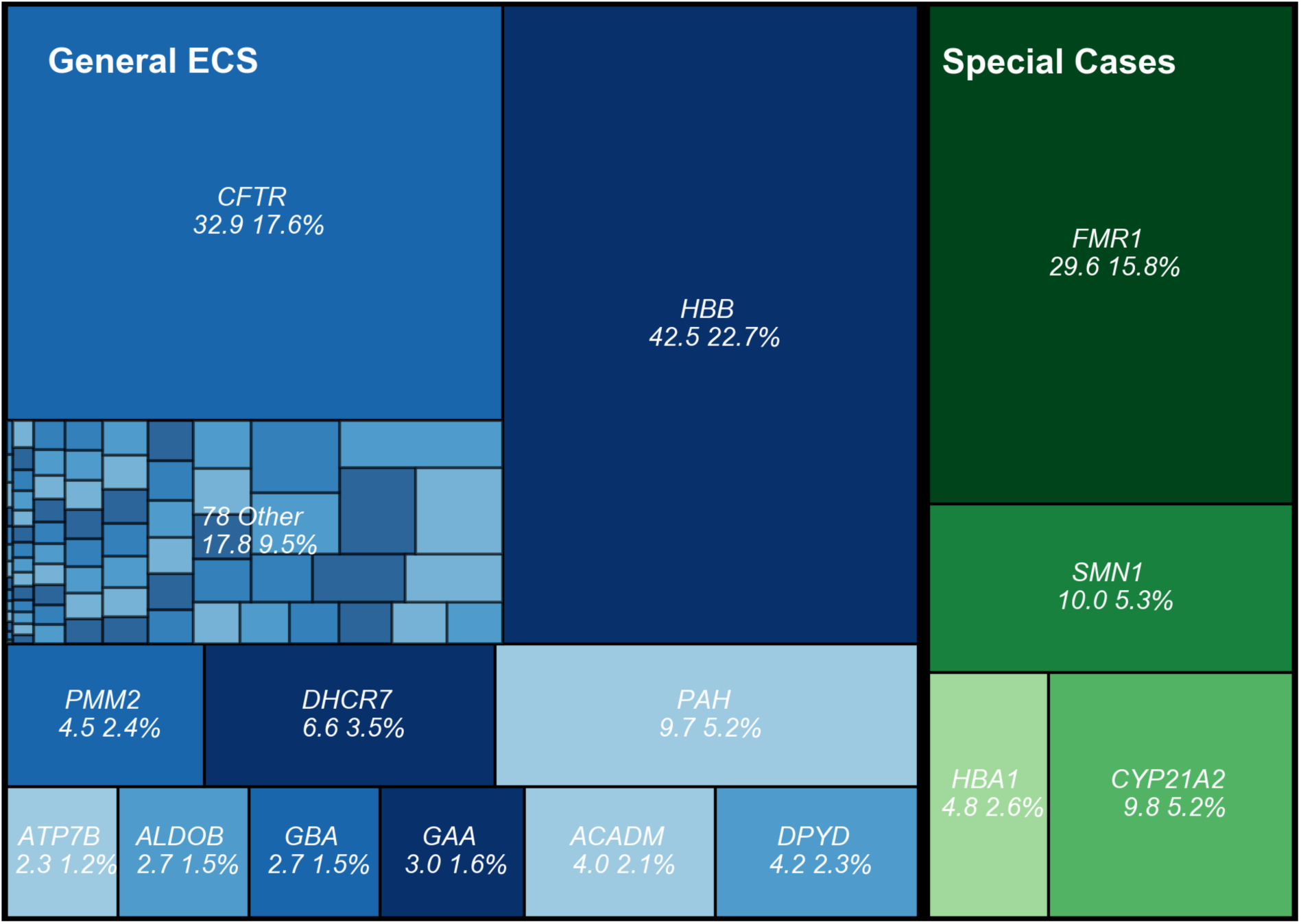
The disease risk contribution of each severe or profound condition on Counsyl Family Prep Screen is shown for a US-census weighted population. 78 conditions contributing fewer than 2 affected fetuses per 100,000 are lumped into one category (“78 Other”) for visual clarity; individual component diseases are outlined but not labeled. For each condition, the number of affected fetuses (per 100,000) is shown, along with the percentage of the total disease risk. The area of each box is proportional to the disease risk. Reported numbers include panel-wide deletion predictions when applicable (see Supplementary Methods).

**Table 2.** Several high-prevalence diseases with known technical challenges are listed below.

| **Disease** | **Gene** | **Why is it difficult?** | **What technology can overcome difficulty?** | **OMIM** |
| --- | --- | --- | --- | --- |
| Fragile X Syndrome | *FMR1* (5' UTR) | Low complexity sequence (CGG Repeats) | PCR + Capillary Electrophoresis <sup>26</sup> | #300624 |
| Congenital Adrenal Hyperplasia | *CYP21A2* | 99% Identical Pseudogene *CYP21A1P* | NGS + Custom Caller <sup>27</sup> | #201910 |
| Alpha Thalassemia | *HBA1/2* | Sequence Identity of *HBA1* and *HBA2* | NGS + Custom Caller <sup>29</sup> | #604131 |
| Gaucher's Disease | *GBA* | 95% Identical Pseudogene *GBAP1* | NGS + Custom Caller <sup>30</sup> | #230800 |
| Spinal Muscular Atrophy | *SMN1/2* | Nearly identical genes (except for 1 base) | qPCR or allele-specific NGS <sup>31</sup> | #253300 |

To further quantify how inclusion of technically challenging variants impacts aggregate panel sensitivity, we constructed three hypothetical full-exon sequencing ECS panels (Figure 2; top bar) for severe and profound diseases. The baseline panel only calls SNPs, indels, and select high-prevalence CNVs (e.g., founder mutations) and excludes several technically challenging diseases (fragile X syndrome, 21-hydroxylase-deficient congenital adrenal hyperplasia, alpha thalassemia, and spinal muscular atrophy; see Discussion and Table 2). The next panel illustrates the marginal gains from adding the technically challenging diseases: excluding these conditions causes 28.9% of affected fetuses to be missed. Finally, the last panel of the top row (Figure 2) assesses the marginal gain of panel-wide exon-level deletion calling (i.e., detection of novel copy-number losses other than known recurrent founder variants). Novel deletion calling adds approximately 4 affected fetuses per 100,000 to the overall detected risk, a relative improvement of 2%. For context, this contribution is roughly comparable to the net contribution of the 50 lowest-prevalence diseases on the panel. The value of novel deletion calling varies by ethnicity: for some ethnicities and diseases, panel-wide deletion calling contributes up to 13 affected fetuses per 100,000 (Supplementary Figures S2, S3).

**Figure 2.**
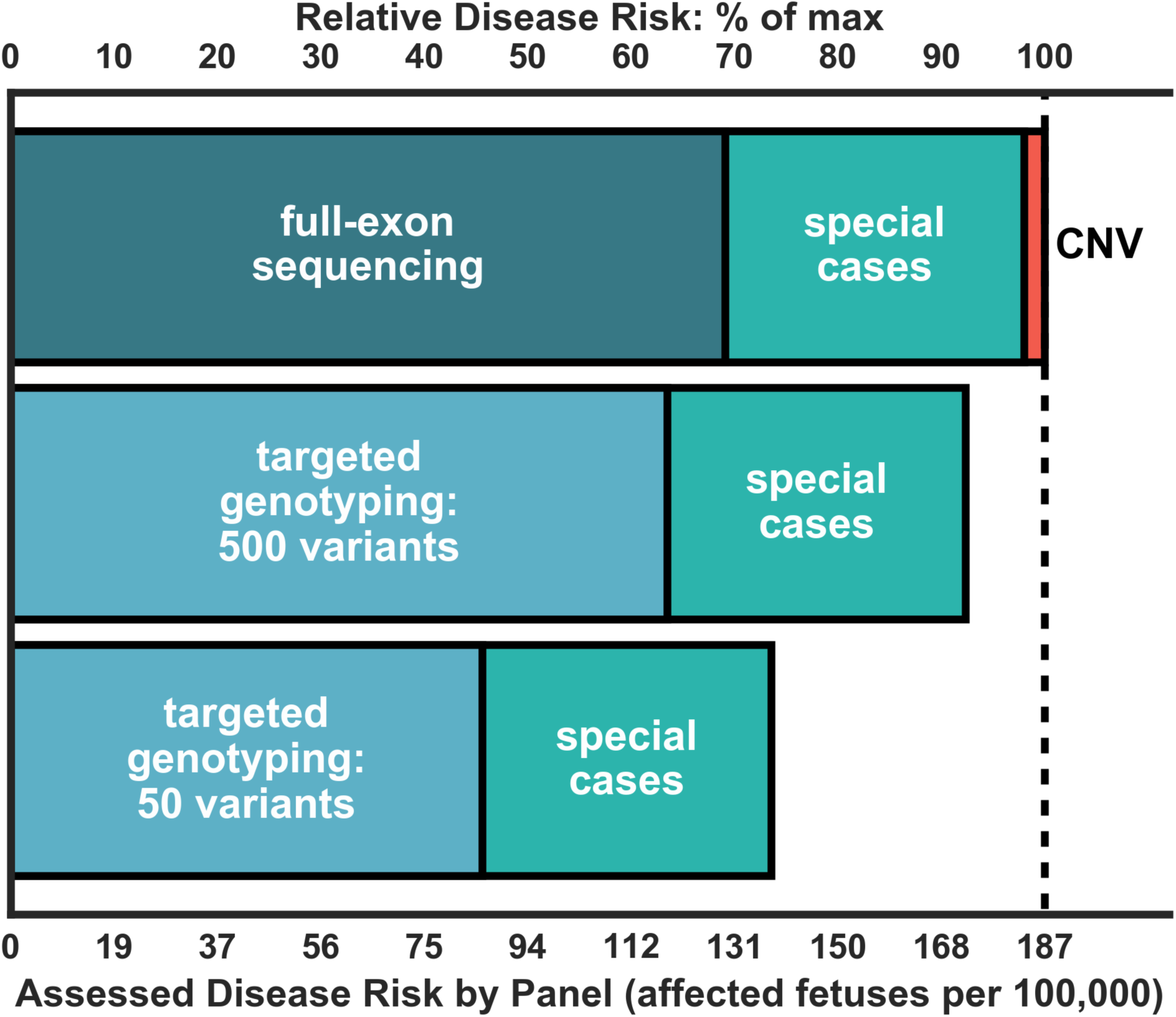
The sensitivity of several hypothetical ECS panels is compared using the disease risk as a proxy. The absolute disease risk, in affected fetuses per 100,000, is plotted on the bottom axis. The top axis shows the contribution as a percent of the total assessed disease risk of the most comprehensive panel considered (full-exon sequencing + special cases + CNV).

To quantify the extent to which it is possible to assess disease risk by targeted genotyping (Figure 2; bottom bars), we constructed sensitivity-optimizing hypothetical targeted genotyping assays of different sizes by selecting from a list of variants rank-ordered by their contribution to overall disease risk (see Supplementary Methods). The targeted genotyping assays detect approximately half the maximal disease risk when technically challenging diseases are omitted. A panel consisting of 500 optimally selected variants plus challenging diseases is capable of detecting 92.4% of the total risk. A per-variant analysis of optimal ECS panels is given in Supplementary Figure S4.

To further clarify how sequencing improves risk assessment at the level of individual diseases, we also estimated the per-gene contribution to disease risk for a previous, targeted genotyping version of Family Prep Screen (Supplementary Table S4). Excluding special cases, the targeted genotyping panel detects 95 affected fetuses per 100,000 births; this number lies between the best-case TG panels of 50 and 500 variants, consistent with the panel probing 332 variants. As expected, some of the conditions with a long history of screening are well captured by the TG panel (CFTR: 1.1 fold gain, HBB: 1.08 fold gain), while other conditions show larger gains for NGS. In aggregate, using NGS leads to a 1.4 fold increase in disease risk as compared to probing the same diseases with the TG panel. Furthermore, the relative gain of NGS (versus TG) may be greater for ethnicities that are poorly served by existing TG panels, such as cystic fibrosis in Asians (Supplementary Fig. S5).

### Real-time curation enables full-exon sequencing

While sequencing maximizes analytic sensitivity (relative to genotyping of selected sites), maintaining high clinical sensitivity and specificity also requires accurate variant interpretation. To assess the sensitivity and specificity of real-time ECS variant interpretation, we compared 505 variant classifications using in-house curation, VEP automated curation, and consensus ClinVar submissions (see Methods). The consensus ClinVar submissions were treated as the reference, though we note that consensus among ClinVar submissions does not necessarily reflect biological truth.

While fully automated computational approaches are much less laborious than full literature-based curation, comparison to ClinVar consensus shows that VEP-based methods must sacrifice either sensitivity or specificity, neither of which is desirable in a screening context (Table 3). In contrast, the integrated interpretation pipeline described achieved both high sensitivity (91.1%) and specificity (99.7%) relative to ClinVar consensus, underscoring the clinical value of combining automated and manual curation.

**Table 3.** Real-time variant curation is evaluated using ClinVar consensus as a reference standard. Two simple VEP curation models are also considered for the sake of comparison. Note that the sensitivity and specificity numbers in this table refer to the variant interpretation process alone, as opposed to the clinical sensitivity (with disease risk as a proxy) and specificity discussed elsewhere in this work. Furthermore, the ClinVar consensus is treated as a gold-standard; sensitivity and specificity here are thus interpreted with respect to concordance to a (possibly imperfect) reference.

| **Curation Method** | **Curation Type** | **Gold Standard** | **Interpretive Sensitivity** | **Interpretive Specificity** |
| --- | --- | --- | --- | --- |
| Counsyl | Manually-Reviewed | ClinVar Consensus | 91.1% | 99.7% |
| VEP Specific | Automated | ClinVar Consensus | 34.6% | 99.7% |
| VEP Sensitive | Automated | ClinVar Consensus | 95.5% | 40.8% |

## DISCUSSION

### Evaluating Expanded Carrier Screening Tests

There is not yet consensus on the “ideal” ECS design. Here, we describe a method in which candidate diseases of high severity are enumerated and then selected for inclusion on a panel based on maximizing aggregate and per-disease sensitivity and specificity. Critically, the optimization of disease-risk sensitivity and specificity differs from a panel-design strategy that simply maximizes the number of genes. Indeed, we argue that disease-risk sensitivity and specificity are better metrics for ECS panel comparisons than gene number, since the latter does not account for features that vary widely across genes, such as disease incidence, assay difficulty, and inheritance pattern.

Assay technology and variant interpretation are key determinants in maximizing ECS sensitivity and specificity. We demonstrate that full-exon sequencing—especially when coupled with technically challenging disease assays and novel CNV detection—provides a sensitivity gain over targeted genotyping. Importantly, these gains require rapid and accurate variant interpretation, which we show can be performed in real-time for novel variants in a manner that is consistent across laboratories.

Building an ECS panel requires careful attention to not only what genes to include, but also what genes to exclude. Sequencing, curating, reporting, and counseling on diseases with limited clinical severity may increase health-care costs and heighten patient anxiety without a commensurate improvement in clinical utility. For these reasons, we have advocated the use of a systematic method for assessing disease severity. Although there are limitations and exceptions to such an approach, we argue that it is a useful starting point for comparisons of ECS panels.

While not the main focus of this work, the cost of performing ECS in the clinic remains a key factor when designing an ECS panel. The ECS panel described herein relies almost exclusively on NGS. With a very low cost per base, NGS coupled with appropriate analysis software and an efficient variant interpretation protocol flexibly accommodates panel updates and can affordably assess carrier status even for technically challenging diseases like alpha-thalassemia and CAH, which historically involve more expensive single-gene assays (e.g., Multiplex Ligation-Dependent Probe Amplification).

The process of ECS panel design should also consider how screening for each disease gene could impact the reproductive options and intervention strategies that patients pursue. For instance, early diet interventions can mitigate the effects of medium chain acyl-CoA dehydrogenase deficiency and phenylalanine hydroxylase deficiency, making them attractive targets for inclusion in ECS. Similarly, early educational interventions can benefit fragile X syndrome patients ^3^, underscoring the value of prenatal screening for this condition. Irrespective of whether patients pursue alternative courses of action in response to ECS, the information and autonomy gained from testing holds value.

### Technically Challenging Genes

Several diseases of high clinical importance (Table 2) have low sequence complexity (e.g., CGG repeats) and/or high homology (e.g., pseudogenes) ^25^, making them difficult to assess with targeted genotyping or full-exon sequencing. These typically require either specialized molecular assays (e.g., testing for fragile X syndrome often uses PCR with capillary electrophoresis ^26^), or custom NGS software (e.g., CAH ^27^). As shown in Figures 1 and 2, these “special case” genes contribute substantially to the population disease risk and in turn, ECS test sensitivity. For example, in order to achieve the same sensitivity, a laboratory could either introduce a complex assay for CAH with 95% sensitivity (9.8 affected births per 100,000) or add the 68 rarest conditions (10.0 affected births per 100,000). This comparison highlights an important feature of panel design: panel constitution and panel size both impact assessed disease risk, and the panel with highest assessed disease risk (and therefore clinical value) may not screen the most genes. Therefore, it is important to quantify each gene’s relative impact on the aggregate assessed disease risk of an ECS.

### Considerations beyond Aggregate Sensitivity and Specificity

We have suggested using the panel-wide sensitivity and specificity as quantitative comparators of ECS panels. In addition, we suggest that per-disease negative predictive value (NPV) is another critical quantity to consider. As a thought experiment, consider two ECS panels that assess the same aggregate disease risk, say, 100 affected pregnancies per 100,000. Suppose the first panel (panel A) screens 100 diseases with 100% sensitivity, while the second panel (panel B) screens 1000 genes with 10% sensitivity. Because their aggregate sensitivities are the same, both panels leave couples with the same overall residual risk of an affected pregnancy after a single-round of ECS. However, they differ in the disease-level NPV. We argue that panel A is preferable because couples receiving negative results for panel A can be confident that they (1) are not at risk for the diseases listed on the panel and (2) are only at risk for diseases conspicuously omitted from the panel.

The argument for prioritizing per-disease NPV provides additional support for the inclusion of panel-wide CNV calling and other sensitivity-boosting improvements to a panel’s existing diseases. Recall that the addition of panel-wide large deletion calling increased the captured disease risk by 2%, with each disease receiving a boost in sensitivity. The net sensitivity gain of deletion calling is roughly equivalent to the addition of approximately 50 new genes. If a panel could add CNV calling or 50 additional genes, we argue that the CNV calling is preferable in order to maximize the panel’s disease-level NPV.

### Application to future panel design

The present work uses retrospective analysis of a commercial ECS to evaluate recommendations for panel design. These recommendations take the form of both general principles (e.g., maximize the sensitivity, specificity, and per-disease NPV for severe and profound diseases) as well as methodological details (e.g., inclusion of hard-to-sequence genes, use of full-exon sequencing, and calling of novel CNVs).

For future panel design, one will likely not have access to 400,000 test results with clinical-grade variant curations. However, disease incidence data will be available, as well as allele frequency data from, e.g., the Exome Aggregation Consortium ^28^. These data could be used to drive a similar analysis in a prospective setting. Care will need to be taken due to unknown variant significance, uncertainty in incidence estimates, and unknown assay detection rate. Despite these challenges, we expect the same principles for optimal ECS panel design to hold.

### Comparison to ACOG Opinion

ACOG very recently released a new opinion ^32^ about carrier-screening panels, making specific recommendations regarding the severity and frequency of diseases that should be screened. Regarding severity, the opinion suggested that screened diseases “have a well-defined phenotype, have a detrimental effect on quality of life, cause cognitive or physical impairment, require surgical or medical intervention, or have an onset early in life.” These criteria are captured well by our disease-severity classification scheme ^2^, summarized in Table 1. With respect to frequency, the opinion supports screening for diseases with a carrier frequency of 1 in 100 or greater. This carrier frequency threshold roughly corresponds to a disease risk of 2.5 per 100,000 births, which is exceeded by 14 conditions in Figure 1. Although our panel does screen some less frequent conditions, we reiterate that the 78 least prevalent conditions (Figure 1) satisfy the recommended severity criteria and have a combined impact of 17.8 per 100,000 births (10% of total risk), sufficiently in excess of the frequency criteria that we favor their inclusion in ECS. Furthermore, several of the 78 least prevalent conditions do meet the 2.5 per 100,000 threshold when considered on a per-ethnicity basis.

## CONCLUSION

We have described a principled method for ECS panel design that selects candidate diseases using systematic severity classification and maximizes sensitivity among those candidates. Based on laboratory testing data, technically challenging genes and full-exon coverage were identified as dominant contributors to overall sensitivity. Furthermore, we argue that the per-disease negative predictive value is a crucial secondary consideration. Clear principles and methodical panel construction ensure transparency in panel design and addresses concerns put forth in guidelines from medical organizations. Broader adoption of these or similar methods would result in consistent panel design and establish a clear basis for panel comparison and evaluation by interested health-care providers.

## Supplementary Information

Supplementary information is available online as a separate PDF.

## REFERENCES

1. Altshuler, D., Daly, M. J. & Lander, E. S. Genetic mapping in human disease. Science 322, 881–888 (2008).

2. Lazarin, G. A. et al. Systematic Classification of Disease Severity for Evaluation of Expanded Carrier Screening Panels. PLoS One 9, e114391 (2014).

3. Hagerman, R. J. et al. Advances in the treatment of fragile X syndrome. Pediatrics 123, 378–390 (2009).

4. American College of Medical Genetics Newborn Screening Expert Group. Newborn screening: toward a uniform screening panel and system--executive summary. Pediatrics 117, S296–307 (2006).

5. Commentary, C. Expanded Carrier Screening in Reproductive Medicine—Points to Consider. doi:10.1097/AOG.0000000000000666

6. American College of Obstetricians and Gynecologists Committee on Genetics. ACOG Committee Opinion No. 486: Update on carrier screening for cystic fibrosis. Obstet. Gynecol. 117, 1028–1031 (2011).

7. Watson, M. S. et al. Cystic fibrosis population carrier screening: 2004 revision of American College of Medical Genetics mutation panel. Genet. Med. 6, 387–391 (2004).

8. Prior, T. W. & Professional Practice and Guidelines Committee. Carrier screening for spinal muscular atrophy. Genet. Med. 10, 840–842 (2008).

9. Lazarin, G. A. et al. An empirical estimate of carrier frequencies for 400+ causal Mendelian variants: results from an ethnically diverse clinical sample of 23,453 individuals. Genet. Med. 15, 178–186 (2013).

10. Azimi, M.et al. Carrier screening by next-generation sequencing: health benefits and cost effectiveness. Mol Genet Genomic Med 4, 292–302 (2016).

11. Grody, W. W. et al. ACMG position statement on prenatal/preconception expanded carrier screening. Genet. Med. 15, 482–483 (2013).

12. Henneman, L.et al. Responsible implementation of expanded carrier screening. Eur. J. Hum. Genet. 24, e1–e12 (2016).

13. Ghiossi, C.et al. Clinical Utility of Expanded Carrier Screening: Reproductive Behaviors of At-Risk Couples. bioRxiv (2016). doi:10.1101/069393

14. Richards, S.et al. Standards and guidelines for the interpretation of sequence variants: a joint consensus recommendation of the American College of Medical Genetics and Genomics and the Association for Molecular Pathology. Genet. Med. 17, 405–424 (2015).

15. Haque, I. S. et al. Modeled Fetal Risk of Genetic Diseases Identified by Expanded Carrier Screening. JAMA 316, 734–742 (2016).

16. US Census. Available at: http://factfinder.census.gov/faces/nav/jsf/pages/index.xhtml. (Accessed: 26th January 2016)

17. Lakeman, P.et al. Three-month follow-up of Western and non-Western participants in a study on preconceptional ancestry-based carrier couple screening for cystic fibrosis and hemoglobinopathies in the Netherlands. Genet. Med. 10, 820–830 (2008).

18. Gross, S. J., Pletcher, B. A., Monaghan, K. G. & Professional Practice and Guidelines Committee. Carrier screening in individuals of Ashkenazi Jewish descent. Genet. Med. 10, 54–56 (2008).

19. Yrigollen, C. M. et al. AGG interruptions within the maternal FMR1 gene reduce the risk of offspring with fragile X syndrome. Genet. Med. 14, 729–736 (2012).

20. Adams, M. C., Evans, J. P., Henderson, G. E. & Berg, J. S. The promise and peril of genomic screening in the general population. Genet. Med. 18, 593–599 (2016).

21. Landrum, M. J. et al. ClinVar: public archive of interpretations of clinically relevant variants. Nucleic Acids Res. 44, D862–8 (2016).

22. McLaren, W.et al. The Ensembl Variant Effect Predictor. Genome Biol. 17, 122 (2016).

23. Nikita Mehta, M. S. et al. Design of scalable gene panels for carrier screening. Presented at ACMG annual meeting. (2016). http://research.counsyl.com/posters/2016/ACMG/ACMG16-Panel-Design.pdf

24. Edwards, J. G. et al. Expanded carrier screening in reproductive medicine-points to consider: a joint statement of the American College of Medical Genetics and Genomics, American College of Obstetricians and Gynecologists, National Society of Genetic Counselors, Perinatal Quality Foundation, and Society for Maternal-Fetal Medicine. Obstet. Gynecol. 125, 653–662 (2015).

25. Mandelker, D.et al. Navigating highly homologous genes in a molecular diagnostic setting: a resource for clinical next-generation sequencing. Genet. Med. (2016). doi:10.1038/gim.2016.58

26. Chen, L.et al. An information-rich CGG repeat primed PCR that detects the full range of fragile X expanded alleles and minimizes the need for southern blot analysis. J. Mol. Diagn. 12, 589–600 (2010).

27. Muzzey, D. An NGS-based Carrier Screen for Congenital Adrenal Hyperplasia with 95% Detection Rate. Presented at ASHG annual meeting. (2015)

28. Lek, M.et al. Analysis of protein-coding genetic variation in 60,706 humans. Nature 536, 285–291 (2016).

29. Maguire JR. D’Auria KM, Lai HH, Wang X, Chu CS, Haque IS, Evans EA, Kang HP, Muzzey D. Next-generation sequencing carrier screen for Alpha Thalassemia identifies both common and rare variants. Presented at AMP annual meeting. (2015). http://research.counsyl.com/posters/2015/AMP/AMPAlphaThal.pdf

30. D’Auria KM, Theilmann MR, Iori K, Chu CS, Haque IS, Evans EA, Kang HP, Maguire JR, Muzzey D. NGS-based carrier screen for Gaucher’s disease calls variants and detects large rearrangements between GBA and GBAP1. Presented at AMP annual meeting (2015). http://research.counsyl.com/posters/2015/AMP/AMPGBA.pdf

31. Wang, X.et al. Next-generation sequencing assay accurately determines carrier status for spinal muscular atrophy. Presented at ASHG annual meeting (2015). http://research.counsyl.com/posters/2015/AMP/AMPSMA.pdf

32. American College of Obstetricians and Gynecologists. Carrier screening in the age of genomic medicine. Committee Opinion No. 690. Obstet Gynecol;129:e35–40 (2017)

